# Cebranopadol, a novel long-acting opioid agonist with low abuse liability, to treat opioid use disorder: Preclinical evidence of efficacy

**DOI:** 10.1101/2023.07.21.550008

**Authors:** Veronica Lunerti, Qianwei Shen, Hongwu Li, Federica Benvenuti, Laura Soverchia, Rajesh Narendran, Friedbert Weiss, Nazzareno Cannella, Roberto Ciccocioppo

## Abstract

The gold standard pharmacological treatment for opioid use disorder (OUD) consists of maintenance therapy with long-acting opioid agonists such as buprenorphine and methadone. Despite these compounds having demonstrated substantial efficacy, a significant number of patients do not show optimal therapeutic responses. Moreover, the abuse liability of these medications remains a major concern. Cebranopadol, is a new, long-acting pan-opioid agonist that also activates the nociception/orphanin FQ NOP receptor. Here we used rats to explore the therapeutic potential of this agent in OUD. First, in operant intravenous self-administration experiments we compared the potential abuse liability of cebranopadol with the prototypical opioid heroin. Under a fixed ratio 1 (FR1) contingency, rats maintained responding for heroin (1, 7, 20, 60 μg/inf) to a larger extent than cebranopadol (0.03, 0.1, 0.3, 1.0, 6.0 μg/inf). When the contingency was switched to a progressive ratio (PR) reinforcement schedule, heroin maintained responding at high levels at all except the lowest dose. Conversely, in the cebranopadol groups responding decreased drastically and the break point (BP) did not differ from saline controls. Next, we demonstrated that oral administration of cebranopadol (0, 25, 50 μg/kg) significantly attenuated drug self-administration independent of heroin dose (1, 7, 20, 60 μg/inf). Cebranopadol also reduced the break point for heroin (20 μg/inf). Furthermore, in a heroin self-administration training extinction/reinstatement paradigm, pretreatment with cebranopadol significantly attenuated yohimbine stress-induced reinstatement of drug seeking. Together, these data indicate that cebranopadol has limited abuse liability compared to heroin and is highly efficacious in attenuating opioid self-administration and stress-induced reinstatement, suggesting clinical potential of this compound for OUD treatment.

## Introduction

Heroin abuse and dependence is a global public health problem. Agonist maintenance therapy with long-acting opioids like methadone and buprenorphine is currently the most effective treatment (1–3). For example, strong preclinical and clinical evidence indicates that buprenorphine dose-dependently reduces opioid use and relapse to drug seeking (4–7). The long-term outcomes of buprenorphine therapy at medium and high doses appears to be as effective as methadone for the treatment of heroin dependence (8, 9). However, buprenorphine is associated with milder side effects compared to methadone, including reduced risk of dysphoria, respiratory depression, and tolerance development (10, 11).

Buprenorphine is classically viewed as a mixed opioid MOP partial agonist and DOP/KOP antagonist. However, at higher concentrations it also activates nociceptin opioid-like receptors (NOP) where it acts as a low efficacy agonist (12–16). Activation of MOP receptors is considered a key mechanism through which buprenorphine attenuates opioid consumption and seeking. However, emerging evidence suggests that concomitant activation of NOP receptors by buprenorphine enhances its safety profile (7, 11, 17) and expands its therapeutic potential to other forms of addiction. It has been established, for example, that buprenorphine reduces alcohol drinking via stimulation of NOP and attenuates cocaine consumption through co-activation of MOP and NOP receptors (12, 18). Moreover, preclinical data using place conditioning and self-administration procedures have shown that activation of NOP reduces the rewarding effects of opioids. This points to the possibility that NOP receptor stimulation may be a novel mechanism for treating opioid addiction (19–21).

Recently, a new spiro[cyclohexane-dihydropyrano[3,4-b]indole]-amine structure characterized by balanced affinity and agonistic potency for MOP and NOP receptors has been developed (22). This includes the compound cebranopadol which has a promising PK/PD profile and has been identified as effective for the treatment of pain (23, 24).

Cebranopadol is an agonist at both NOP and MOP receptors. Radioligand binding assays revealed sub-nanomolar affinity for both rat and human NOP and MOP receptors and an additional 20 times lower affinity for DOP and KOP receptors (25). Because of this pharmacodynamic profile and the long-acting pharmacokinetics of the compound, we predicted that this compound would have significant potential as a novel therapeutic for opioid use disorder (OUD).

Abuse liability typically associated with opioid based therapies is a concern. Since cebranopadol is a potent MOP agonist, we sought to evaluate its abuse potential using intravenous self-administration procedures. Additionally, we conducted a series of experiments to explore the effect of cebranopadol on heroin self-administration and yohimbine stress-induced reinstatement of drug seeking in the rat.

## Methods and materials

### Animals

Experiments were performed in male Wistar rats (Charler River Laboratories, Italy) weighing 280– 330g at the beginning of the experiments. Rats were housed two per cage in a room with a reverse 12:12h light/dark cycle (lights off at 8:00 a.m.), constant temperature (20–22°) and humidity (45– 55%). Rats had *ad libitum* access to food (4RF18, Mucedola, Settimo Milanese, Italy) and tap water except during experimental time. Prior to experiments, animals were acclimated to the vivarium for one week and handled daily for 5 min. Experiments were conducted during the dark phase of the light/dark cycle. All procedures were carried out in accordance with the *European Community Council directive for Care and Use of Laboratory Animals* and the *National Institutes of Health guidelines for the care and use of laboratory animals*.

### Drugs

Heroin (NIDA Drug Supply System, Bethesda, USA) was dissolved in sterile physiological saline (0.9%) at the doses of 1, 7, 20, and 60 µg/0.1 ml/infusion.

Cebranopadol (Biochempartner Co., Ltd, China) was dissolved and diluted with 10% DMSO + 5% Cremophor EL + 85% saline. Cebranopadol or its vehicle were administered orally (po) by the experimenter 1h before test sessions (25). In self-administration experiments designed to ascertain the drug’s abuse liability, cebranopadol was self-administered intravenously (i.v.).

Yohimbine hydrochloride (Sigma, Italy) was dissolved in millipore water at a dose of 1.25 mg/kg and administered intraperitoneally (ip) 30 min before tests.

### Self-administration apparatus

Training and testing were conducted in operant self-administration (SA) chambers (Med Associate Inc.), enclosed in sound attenuating, ventilated environmental cubicles. Each chamber was equipped with two retractable levers located on a side panel (4 cm above the grid floor), two stimulus lights above each lever, a house light and two sound generators. An external syringe pump controlled the delivery (0.1 ml) of drug solutions. Responses on the right (active) lever at the required ratio resulted in activation of the pump whereas responses on the left (inactive) lever were recorded but had no programmed consequences. During SA, the infusion pump was connected to a plastic tube which ran through a protective metal coil into the SA chamber where it connected to a catheter implanted in the intra-jugular vein of the rat. MEDPC-IV^®^ windows-compatible software was used to control fluid delivery, presentation of visual and sound stimuli, and data collection.

### Intravenous catheter implantation

Animals were anesthetized by intramuscular injection of 100-150 µl of a solution containing tiletamine chloralhydrate (58.17 mg/ml) and zolazepam chloralhydrate (57.5 mg/ml). For i.v. surgeries, incisions were made to expose the right jugular vein. A catheter made from micro-renathane tubing (I.D = 0.020 inches, O.D = 0.037 inches) was subcutaneously routed from the animal’s back to the jugular vein. After insertion into the vein, the proximal end of the catheter and the vein were anchored to the underlying muscle with surgical silk. The distal end of the catheter was attached to a stainless-steel cannula bent at a 90° angle. The cannula was inserted into a dental cement support on the back of the animals and covered with a plastic cap. Immediately after surgery, rats received 200 µL of enrofloxacin (50 mg/ml, Baytril, Germany) intramuscularly and kept warm until fully awake.

Rats were allowed to recover one week before the beginning of SA training. Catheter patency was confirmed by intravenous injection of 150 µl of pentothal sodium (25 mg/ml, Intervet, Italy). Before each SA session catheters were flushed with 100 µl of heparinized saline (20 U.I./ml) containing 1.0 mg/ml of enrofloxacin.

### Self-administration experiments to assess the abuse liability of cebranopadol and heroin under FR and PR contingencies

#### Training

Rats were first trained to self-administer heroin in 2-hour daily sessions. Lever activation under a fixed ratio 1 (FR1) schedule of reinforcement was reinforced by 20 µg/0.1ml infusion (inf) of heroin. Following each heroin infusion, a 20 sec time out (TO) period was in effect during which responses at the active lever had no programmed consequences. Each heroin infusion was paired with illumination of the cue-light above the active lever, which remained on during the TO period. Additionally, throughout the entire SA session, an intermittent beeping tone was presented.

#### Test

Following heroin SA training, animals were divided into heroin SA and cebranopadol SA cohorts. Heroin cohort doses included 1 µg/inf (N=8), 7 µg/inf (N=9), 20 µg/inf dose (N=10), and 60 µg/inf (N=9). Cebranopadol cohort doses included 0.03 µg/inf (N=9), 0.1 µg/inf (N=8), 0.3 µg/inf (N=8), 1.0 µg/inf (N=9), and 6.0 µg/inf (N=8). A separate group (N=8) was trained in heroin SA and then switched to saline and served as a control for both heroin and cebranopadol SA groups. Cebranopadol, heroin and saline SA was maintained for four consecutive days under FR1. On the fifth day groups were switched to a progressive ratio (PR) schedule of reinforcement. Under the PR schedule, the responses required to receive a single heroin infusion increased according to the following sequence: 1, 2, 4, 6, 9, 12, 15, 20, 25, 32, 40, 50, 62, 77, 95, 118, 145, 178, 219, 268, 304 etc. The PR sessions stopped after 4 hours or if the required ratio was not reached within 1 hour, whichever came first. The breakpoint (BP), corresponding to the last completed ratio was used as a measure of motivation for the drugs. In these experiments, the doses of heroin were chosen based on earlier studies (26–28), whereas cebranopadol doses were identified based on its solubility and pharmacokinetic/pharmacodynamic profile (25).

### Effect of cebranopadol on heroin self-administration

The four groups trained to self-administer four different doses (1.0, 7.0, 20.0, and 60.0 μg/inf) of heroin were used. After completion of the PR test, rats were re-trained for FR1 self-administration of heroin for five additional days to reestablish the SA baseline. The effect of cebranoapdol (0.0, 25.0, 50.0 μg/kg) was then evaluated against each of the four heroin doses. Rats received the two doses of cebranopadol and its vehicle in a Latin-square counterbalanced order. Tests were repeated every third day. On the first intervening day rats remained in their home-cage and on the second they were given a baseline heroin SA session.

### Effect of cebranopadol on motivation for heroin

In a separate group of rats (N=8), we tested the effect of cebranoapdol (0.0, 25.0, 50.0 μg/kg) on motivation for heroin (20.0 μg/inf) expressed by the break-point reached in a PR SA session. Rats received the two doses of cebranopadol and its vehicle in a Latin-square counterbalanced order. PR tests were repeated every third day. On the first intervening day rats remained in their home-cage and on the second they were given a baseline heroin SA session under a FR1 contingency.

### Effect of cebranopadol on yohimbine-induced restatement of heroin seeking

This experiment was conducted with a new group of rats (N=11) and consisted of three phases.

#### Self-administration training

Rats were trained to self-administer heroin (20 μg/infusion) in 2-hour daily sessions as described above.

#### Extinction training

Once rats reached a stable SA baseline, they were switched to a 1-hour daily extinction session for 10 consecutive days. During these sessions, responses at the active lever activated the syringe pump but the heroin solution was not delivered.

#### Yohimbine-induced reinstatement of heroin seeking and cebranopadol treatment

On the test day, cebranopadol (0, 25, 50 μg/kg) was given p.o. 60 minutes before the beginning of the reinstatement testing session. This was followed by an intraperitoneal injection of yohimbine (1.25 mg/kg) 30 minutes before the beginning of the session. Yohimbine tests were repeated every fourth day and cebranopadol doses were administered in counterbalanced order according to a Latin square design. Between tests, rats were subjected to extinction sessions.

### Statistical analysis

In the dose/response experiments, one-way between-subjects ANOVA was used to evaluate differences in baseline heroin (20 µg/inf) infusions before rats were divided into various dosing groups of cebranopadol, heroin or saline. Heroin and cebranopadol self-administration at different doses were separately analyzed by two-way ANOVA with one between factor (infusion dose) and one within factor (time). Breakpoints and relative infusions reached under the PR contingency for cebranopadol and heroin were analyzed separately by one-way ANOVA with drug dose as between subjects factor.

The effect of cebranoapdol on heroin self-administration under FR1 contingency was analyzed by one-way within-subjects ANOVA. Each heroin concentration was analyzed separately. The same statistical approach was applied to analyze the effect of cebranopadol on motivation for heroin.

Yohimbine-induced reinstatement was analyzed by paired t-test, comparing responses during the last day of extinction with yohimbine effects in the cebranopadol vehicle condition. The effect of cebranopadol on yohimbine-induced reinstatement was analyzed by one-way within-subjects ANOVA. Active and inactive lever responses were always analyzed separately.

Newman-Keuls and Dunnet’s tests were used for *post-hoc* analysis when appropriate. Results were expressed as mean ± SEM. Statistical significance was set at *p*< 0.05.

## Results

### Cebranopadol shows lower abuse liability potential compared to heroin

#### FR1 schedule of reinforcement

Following heroin (20μg/inf) training, rats were subdivided into heroin dose/response, cebranopadol dose/response, and saline cohorts. At baseline, the resulting ten experimental groups did not differ in the number heroin infusions [F(9, 77)=1.0; *p = not significant (NS)*].

In the heroin dose/response experiment, when groups were switched to different heroin SA doses or saline, ANOVA revealed a significant effect of dose [F(4, 40)=3.4; *p*<0.05], time [F(3, 120)=6.2; *p*<0.001], and dose by time interaction [F(12, 120)=3.0; *p*<0.01]. The number of infusions was highest at the two lowest doses of 1 μg/inf and 7 μg/inf, which induced comparable levels of self-administration. The number of infusions was lower at 20 μg/inf and 60 μg/inf. Each heroin dose sustained self-administration, while the saline group progressively showed extinction of operant responding over time (**Figure 1A**). When the total amount of heroin taken was analyzed, there was an overall effect of dose [F(3,33)=56.4; *p*<0.0001], but no effect of time [F(3,99)=1.0; *NS*] or dose by time interaction [F(9,99)=1.9; *NS*]. The Newman-Keuls post-hoc test within the main dose effect revealed significant differences (p<0.01 and p<0.05) between the different doses (**Figure 1B**). The largest amount of heroin intake occurred at the highest concentration (60 μg/inf) and progressively decreased with decreasing dose (20 μg/inf, 7 μg/inf and 1 μg/inf). Inactive lever responses were always low and did not differ between groups (dose [F(4,40)=1.3; *NS*], time [F(3,120)=2.5; *NS*], time by dose interaction [F(12,120)=1.3; *NS*]).

**Figure 1.**
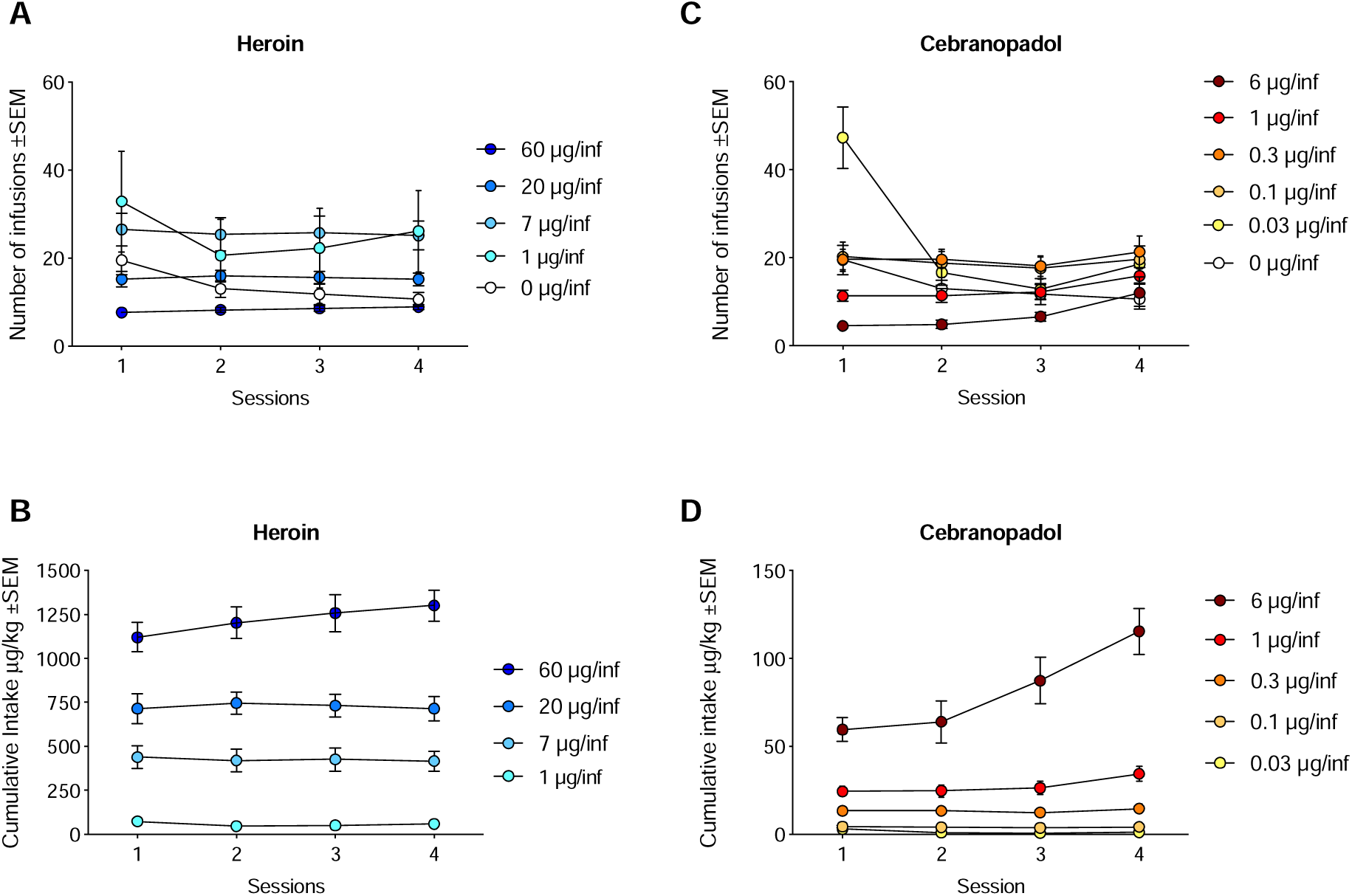
Intravenous FR1 self-administration of different doses of heroin and cebranopadol. **A)** Number of heroin infusions over four consecutive SA days. Data show an inverse relation between heroin dose and number of infusion earned; i.e., the two lowest doses (1.0 and 7.0 µg/inf) were associated the with highest number of infusions. **B)** Cumulative heroin intake. Data indicate that consumption increases as a function of dose. **C)** Number of cebranopadol infusions at different drug doses over four self-administration days. **D)** Cumulative cebranopadol intake. Data demonstrate the highest consumption at the highest dose tested. Data are expressed as Mean ± SEM. Statistically significant differences are reported in the text; stars are omitted for sake of clarity.

When additional heroin trained rats were switched to different doses of cebranopadol or to saline, there was an overall effect of dose [F(5,44)=7.0; *p*<0.0001], time [F(3,132)=16.1; *p*<0.0001] and dose by time interaction [F(15,132)=12.2; *p*<0.0001]. The groups receiving saline or the lowest dose (0.03 µg/inf) of cebranopadol showed an extinction-like behavior as revealed by the progressive decrease in the number of infusions earned over time (days). Larger doses of cebranopadol maintained operant responding over time with the lowest number of infusion at the 6 µg/inf dose (**Figure 1C**). When the total amount of cebranopadol self-administered was analyzed, ANOVA revealed a significant effect of dose [F(4,37)=70.8; *p*<0.0001], time [F(3,111)=10.5; *p*<0.0001], and a dose by time interaction [F(12,111)=7.0; *p*<0.0001]. Newman-Keuls post-hoc tests indicated that the highest dose group (6 µg/inf) showed the largest drug intake compared to the other groups at each time point (**Figure 1D**). Inactive lever responses were always low and did not differ between groups: dose [F(5,44)=6.0; *p*<0.001], time [F(3,1132)=1.5; *NS*], time by dose interaction [F(15,132)=2.7; *NS*].

When rats were tested under the progressive ratio contingency and the break point for heroin was analyzed, ANOVA confirmed a significant overall effect of dose [F(4,40)=7.5; *p*<0.0001]. Newman-Keuls post-hoc test revealed that rats receiving 7, 20 or 60 µg/inf of heroin reached break points significantly higher compared to saline controls (p<0.05 and p<0.001). At the dose of 1 µg/inf heroin failed to maintain operant responding under the PR contingency and the BP was indistinguishable from that of saline controls (**Figure 2A**). ANOVA applied to the number of heroin infusions under PR contingency confirmed a significant overall effect of dose [F(4,40)=19.7; *p*<0.0001]. Post-hoc tests showed a significant (p<0.001) difference between saline and heroin at 7, 20 or 60 µg/inf but not 1 µg/inf (**Figure 2B**).

**Figure 2.**
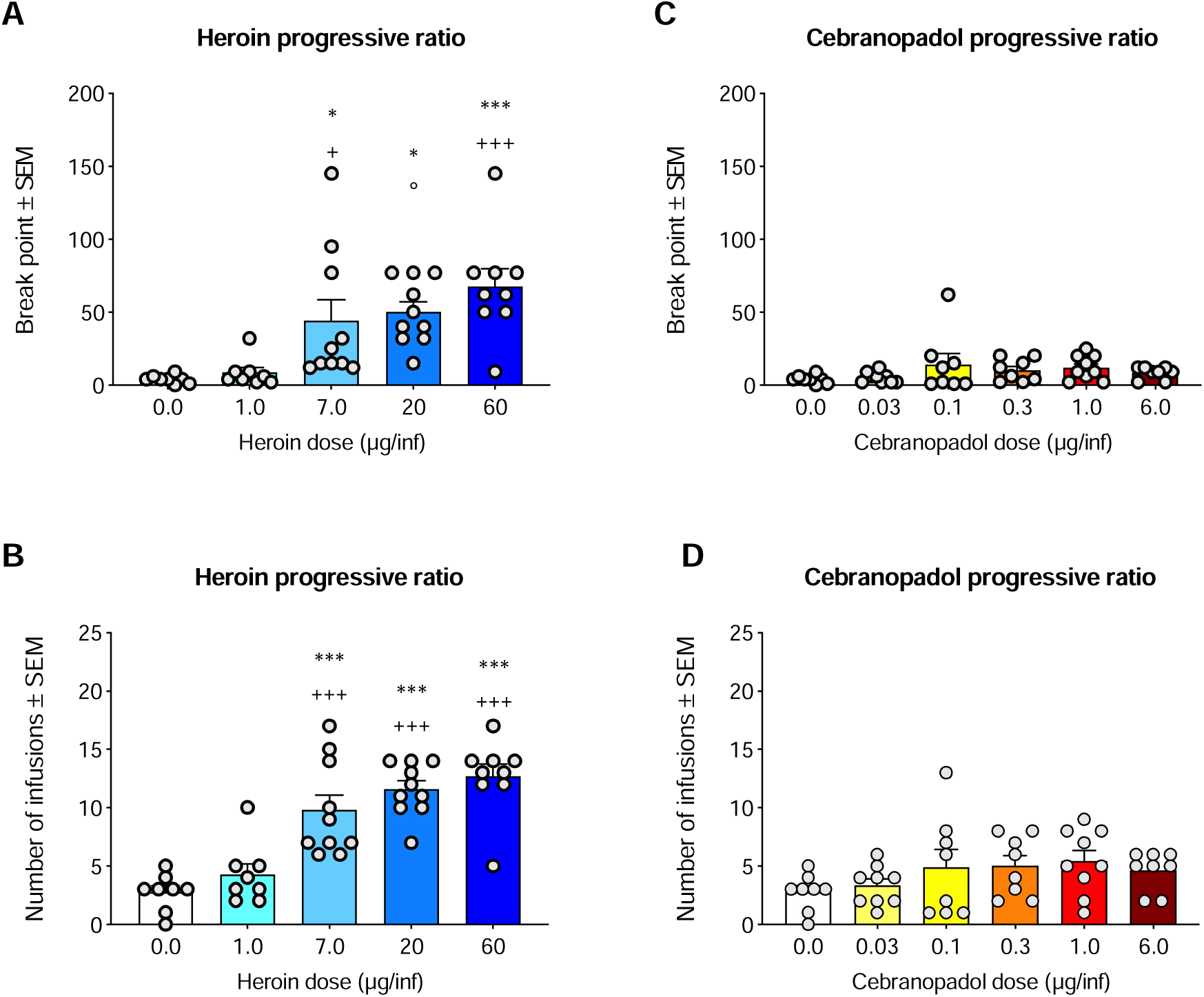
Motivation for different doses of heroin and cebranopadol evaluated on a progressive ratio schedule of reinforcement. **A**) At a dose of 1 µg/inf the break point (BP) for heroin did not differ from saline. The BP progressively increased as a function of dose. **B**) Number of heroin infusions earned during the PR session. **C**) Break points for cebranopadol were very low and did not differ from saline. **D**) Likewise, the number of Cebranopadol infusions earned in the PR session were low and never different from saline. Data are expressed as Mean ± SEM. *p<0.05 and ***p<0.001 from saline; ^+^p<0.05 and ^+++^p<0.001 from heroin 1 µg/inf.

In contrast to heroin, cebranopadol did not show any significant effects either in the BP [F(5,44)=1.3; *NS*] (**Figure 2C**) or the number of infusions [F(5,44)=1.4; *NS*] (**Figure 2D**). None of the cebranopadol groups differed from the saline group.

### Cebranopadol attenuates heroin self-administration

When the effect of cebranopadol on heroin self-administration was analyzed, ANOVA revealed an overall effect of cebranopadol on heroin self-administration at 1 μg/inf [F(2,6)=21.3, *p*< 0.01], 7 μg/inf [F(2,9)=25.2, *p*< 0.0001], 20 μg/inf, [F(2,9)=16.8, *p*<0.001] and 60 μg/inf [F(2,8)=3.7, *p*=0.05]. As shown in **Figure 3A-D**, Dunnet’s post hoc analysis indicated that in the groups receiving 1 μg/inf or 7 μg/inf of heroin, both doses (25 and 50 μg/kg) of cebranopadol markedly (p<0.01 and p<0.001) attenuated drug self-administration. In the group receiving 20 μg/inf of heroin, cebranopadol was also efficacious at 25 μg/kg (p<0.05) and at 50 μg/kg (p<0.001). However, against the highest heroin dose (60 μg/inf) cebranopoadol was efficacious only at 50 μg/kg (p<0.05). Responses at the inactive lever were always very low and not significantly modified by cebranopadol (7 μg/inf, [F(2,9)=1.2, *NS*], 20 μg/inf, [F(2,9)=0.9, *NS*] and 60 μg/inf, [F(2,8)=1.3, *NS*]) except for the 1 μg/inf heroin dose [F(2,6)=7.2, *p*<0.05], where the low number of inactive lever presses (4 ± 1.4) was reduced by both doses of cebranopadol (cebranopadol 25µg/kg, 0.3 ± 0.3; cebranopadol 50µg/kg, 0).

**Figure 3.**
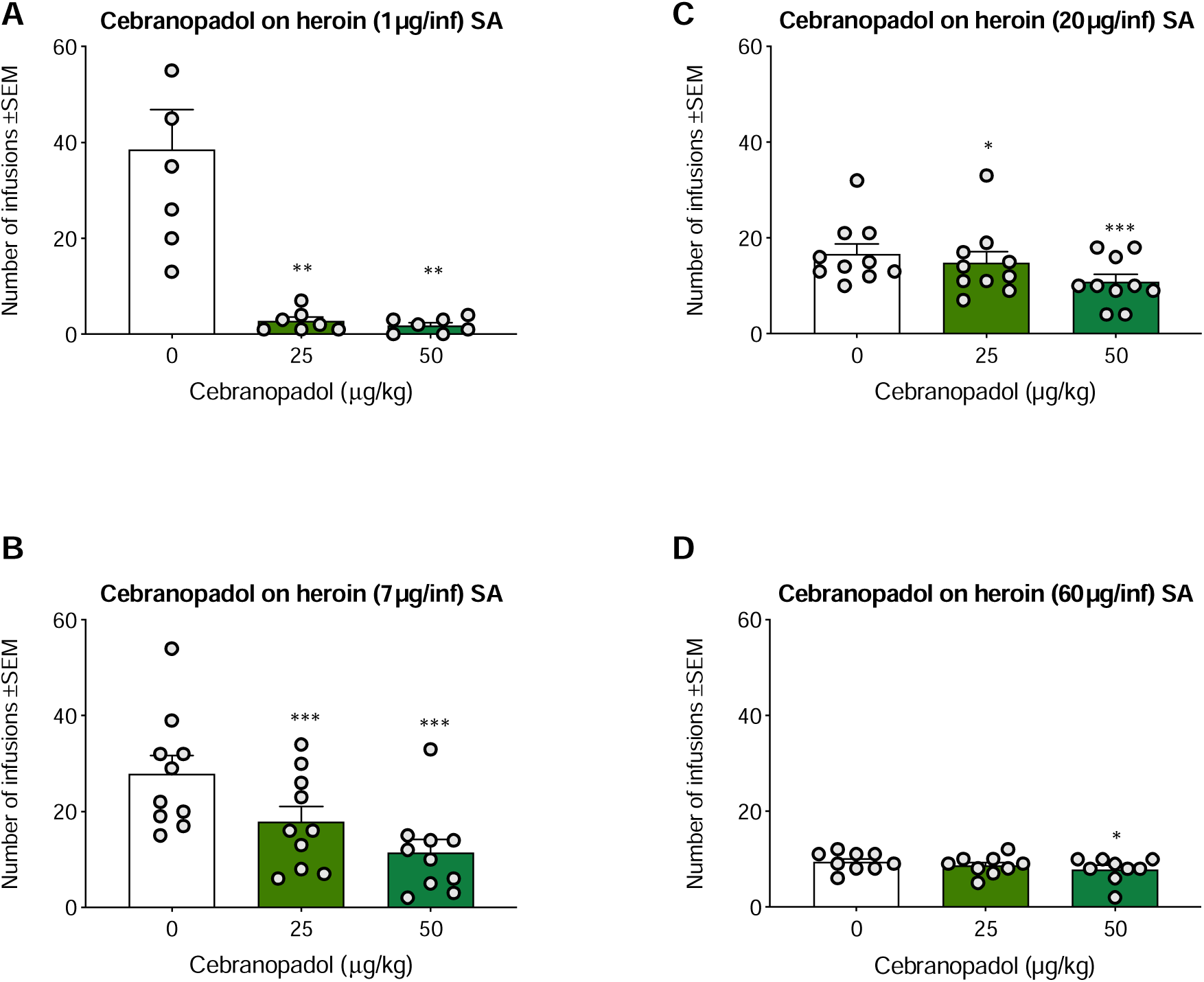
Effect of cebranopadol on FR1 SA of different doses of heroin. Both doses of cebranopadol decreased the number of infusions of **A**): 1 µg; **B**) 7 µg and **C**) 20 µg of heroin. **D)** At 60 µg/inf). Heroin consumption was significantly attenuated only after administration of 50 µg/kg of cebranopadol. Data are expressed as Mean ± SEM. *p<0.05, **p<0.01, and ***p<0.001 vs cebranopadol 0.0 µg/kg dose.

### Cebranopadol attenuates motivation for heroin

One rat lost catheter patency before completing the Latin-square design, and therefore statistical analyses were conducted on N=7 rats. To evaluate the effect of cebranopadol on motivation for heroin both break-points and number of infusions were analyzed. ANOVA of break-points established an overall effect of cebranopadol doses [F(2,6)=33.4, *p*<0.001]. Dunnet’s post-hoc analysis revealed that cebranopadol significantly reduced break-points for heroin at both 25 µg/kg (p<0.05) and at 50 µg/kg (p<0.001) doses (**Figure 4A**). Similarly, when the number of infusions was analyzed, ANOVA showed an overall effect of cebranopadol doses [F(2,6)=36.4, *p*<0.0001], Dunnet’s post-hoc analysis revealed that cebranopadol significantly reduced heroin infusions earned under the PR contingency at both 25 µg/kg (p<0.05) and at 50 µg/kg (p<0.001) doses (**Figure 4B**).

**Figure 4.**
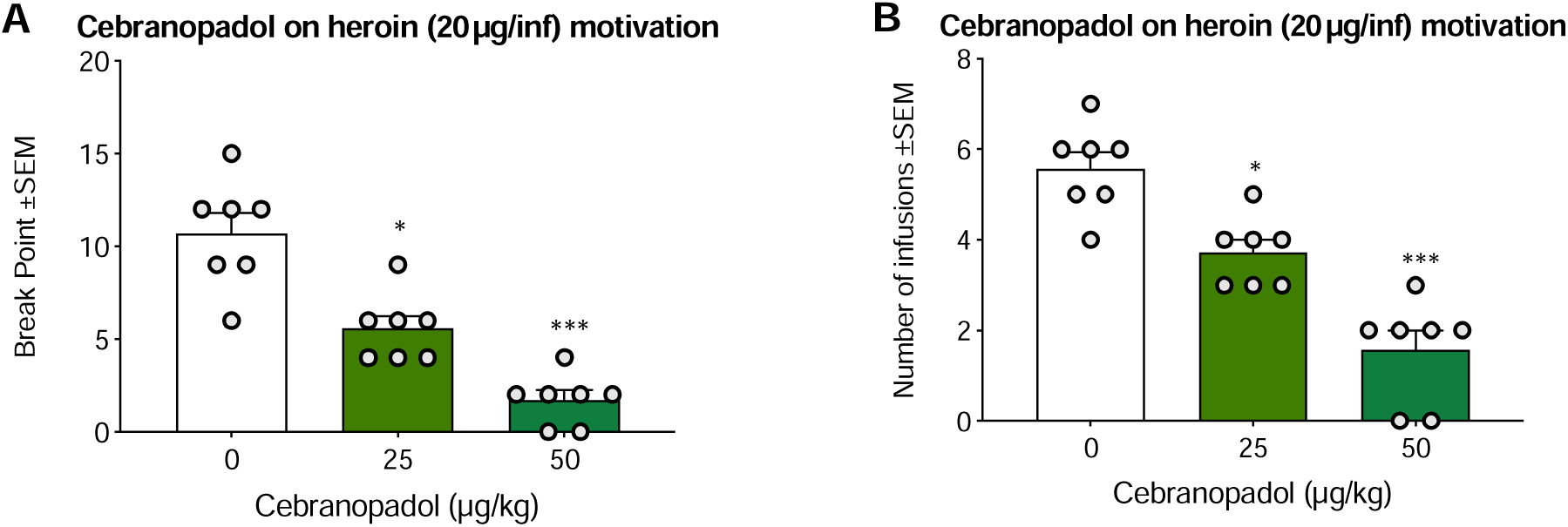
Effect of cebranopadol on the motivation for 20 µg/inf of heroin under progressive ratio schedule of reinforcement. Both doses of cebranopadol decreased: **A**) the break-point for heroin and **B**) the number of infusions earned. Data are expressed as Mean ± SEM. *p<0.05 and ***p<0.001 vs cebranopadol vehicle (0 µg/kg).

### Cebranopadol blocks yohimbine-induced restatement of heroin seeking

One rat was excluded from statistical analysis because of aberrant behavior in response to yohimbine with 1965 lever presses in one hour. At the end of SA training, rats self-administered an average of 15.4±5.2 heroin infusions in the 2-hour session. During extinction, active lever pressing progressively decreased from 47.8±20.05 on the first day to 6.4±1.3 on the last day. When rats received yohimbine, operant responding almost doubled (13.2 ± 2.1) compared to extinction levels and statistical analysis revealed a significant difference [t(9)=2.9; *p*<0.05] between the two conditions. Cebranopadol produced an overall reduction of yohimbine-induced reinstatement [F(2, 9)=3.7, *p*<0.05]. Post-hoc analysis revealed that 50 μg/kg cebranopadol significantly (p<0.05) reduced active lever responses compared to vehicle-treated rats (**Figure 5A**). When inactive lever presses were analyzed, the paired t-test revealed a significant increase in lever responses following yohimbine [t(9)=4.0; *p*<0.01] and ANOVA a significant reduction of lever responses [F(2, 9)=3.8, *p*<0.05] following cebranopadol (**Figure 5B**).

**Figure 5.**
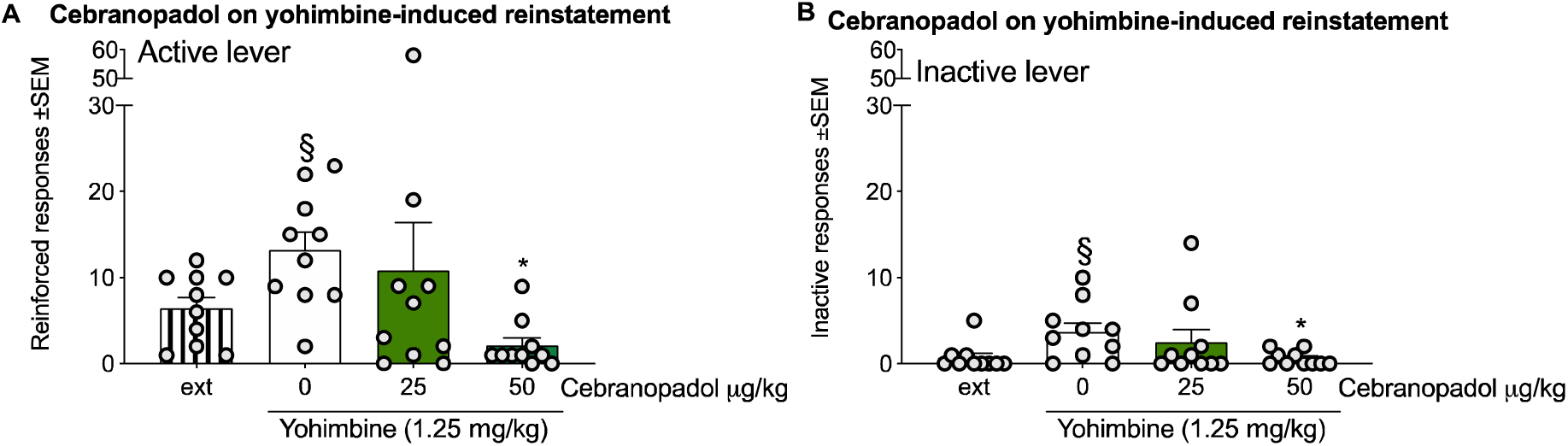
Effect of cebranopadol on yohimbine stress-induced reinstatement of heroin seeking. **A**) Yohimbine produced reinstatement of heroin seeking measured as the increase in active lever responding in cebranopadol vehicle treated rats with respect to extinction (ext). Treatment with 50.0 µg/kg of cebranopadol decreased active lever pressing abolishing heroin seeking. **B**) Responses at inactive lever were also increased by yohimbine and reverted by 50.0 µg/kg of cebranopadol. Data are expressed as Mean ± SEM. ^§^p<0.05 vs extinction (Ext) and *p<0.05 vs cebranopadol 0.0 µg/kg dose.

## Discussion

Cebranopadol is a novel compound in advanced clinical development with comparable subnanomolar binding affinities and full agonist potencies at both NOP and MOP receptors (24, 25). Compared with the prototypic opioid receptor agonist morphine, cebranopadol showed 180∼4800 times higher potency in a variety of animal pain models (25). Moreover, recent studies by our and other laboratories have documented efficacy of cebranopadol for reducing cocaine self-administration and relapse in the rat (29, 30). Notably, conditioned place preference data in rodents and initial clinical observations suggested that, compared to classical opioid agonists, cebranopadol has fewer side effects, such as alteration in motor coordination, depression of respiratory function, tolerance to its analgesic effects, and lower rewarding effects (25, 30–33). A dominant hypothesis in the field is that this favorable pharmacological profile results from the ability of cebranopadol and similar molecules to co-activate NOP and MOP receptors (34, 35).

Abuse liability is a critical side effect of opioid agonists that significantly hampers their therapeutic potential and is a major concern for the use of these agents as analgesics and medications for drug dependence. Here, we conducted an in-depth analysis of cebranopadol’s abuse liability potential compared to the classical opioid agonist heroin (36). Animal pain models indicated that cebranopadol has an EC_50_ of 5.6 μg/kg when given i.v. and 25.1 μg/kg following oral administration (25). Based on these data, in i.v. self-administration experiments we used doses of 0.03, 0.1, 0.3, 1, μg/inf and a maximal dose of 6 μg/inf. At this dose range, cebranopadol is pharmacologically efficacious and soluble in an i.v. injectable solution. At the same time, it does not saturate opioid receptors, allowing for the maintenance of operant self-administration. Heroin, which has a lower affinity for MOP compared to cebranopadol, was given at a higher dose range (1-60 μg/inf) which is known to maintain operant self-administration (26–28). As expected, heroin supported FR-1 lever pressing in a dose-dependent manner, with 7 μg/inf producing the highest levels of responding whereas the highest dose of 60 μg/inf produced the lowest levels of responding. When tested under the PR contingency at 1 μg/inf, heroin did not maintain operant responding, suggesting that at this dose the drug is not reinforcing. Conversely, the PR was maintained, and high BPs were reached when heroin was given at higher concentrations. Cebranopadol also supported FR1 self-administration, indicating that the compound has positive reinforcing effects. Similar to heroin, the number of responses was highest at 0.3 μg/inf whereas the maximum amount of drug taken occurred at the highest concentration (6 μg/inf). With the exception of the lowest dose, the number of cebranopadol infusions did not change over days indicating that maximal reinforcing effect of the drug was reached with the first drug experience and did not increase over time. The effects of cebranopadol and heroin were very different when these drugs were tested under the PR contingency. Heroin supported a very high PR reflecting strong reinforcing actions. The BP was high for each heroin dose except for 1 µg/inf, with a trend toward a higher BP at the highest concentration (60 μg/inf). In contrast, the BP for cebranopadol was very low, not affected by the infusion dose, and always similar to that in saline controls. These findings corroborate initial place conditioning studies in rodents and self-administration experiments in monkeys showing that cebranopadol can have positive reinforcing effects that are, however, much weaker than those of other opioid agonists (29, 30, 37).

Considering the high affinity and potency of cebranopadol for MOP receptors, the degree of abuse potential predicted by our results appears surprisingly low (Linz *et al*, 2014; Linz *et al*, 2017). On the other hand, this finding corroborates the hypothesis that molecules that coactivate MOP and NOP receptors may have a favorable abuse liability profile compared to agonists at classical opioid receptors. The low abuse potential of cebranopadol prompted us to expand our investigation to evaluate its efficacy as a possible treatment for OUD. We therefore tested the effects of cebranopadol on heroin self-administration and reinstatement of heroin seeking induced by yohimbine stress. The results showed that cebranopadol markedly reduced heroin intake at low (1-7 μg/inf) or intermediate (20 μg/inf) doses. However, cebranopadol was less efficacious against the highest (60 μg/inf) dose of heroin. This finding replicates previous observations in human laboratory studies in which the effects of buprenorphine were tested against self-administration of different intranasal heroin doses (38). These earlier studies, in fact, demonstrated that buprenorphine, a compound that shows some similarities with cebranopadol, can attenuate the self-administration of low (12.5 mg) and intermediate (25-50 mg) but not high (100 mg) doses of heroin. Based on the estimated dissociation constant and efficacy, those authors concluded that in order to block the reinforcing effects of high heroin doses it is necessary to inactivate (occupy) more than 90% of available MOP receptors, a criterion that was not achieved by buprenorphine in their study (38). Similarly, it is possible that the doses of cebranopadol used in our experiments may not have produced a sufficiently high level of MOP receptor occupancy, allowing the highest dose of heroin (60 μg/inf) to surmount the effect of the drug. On the other hand, this does not preclude clinical potential of this drug for OUD treatment. In fact, because of its good tolerability and moderate abuse liability, cebranopadol can likely be administered at doses that can produce full MOP receptor occupancy.

One of the major challenges for the treatment of OUD is the high rate of relapse after abstinence. Stress is a major factors triggering drug seeking and relapse (39). Clinical evidence exists that administration of the pharmacological stressor yohimbine, an alpha-2-adrenoceptor antagonist, elicits marked increases in opioid craving in dependent individuals (40). Similar to humans, yohimbine elicits drug seeking in laboratory animals trained to self-administer heroin, which provides translational validity to the use of yohimbine as a model of stress (40–43).

Based on these considerations, in order to further explore the therapeutic potential of this compound, we sought to evaluate the efficacy of cebranopadol in protecting rats from reinstatement of drug seeking elicited by yohimbine. Consistent with published data (41, 42), yohimbine elicited significant reinstatement of heroin seeking. Cebranopadol significantly attenuated this behavior at the highest dose (50 μg/kg). In addition to producing reinstatement at the active lever, yohimbine also elicited an increase in responding at the inactive lever, that can be interpreted as a general arousal phenomenon and/or a strong generalized attempt by the animal to receive heroin. We found that cebranopadol also reduced inactive lever responses in both reinstatement and heroin 1µg/inf self-administration experiments, which possibly could raise issues concerning the specificity of its action. On the other hand, the fact that inactive lever responses were not altered in the self-administration experiments with the three more reinforcing doses of heroin (7-60µg/kg), together with published data showing that cebranopadol did not affect operant responding for saccharin and sweetened condensed milk (29, 30) would seem to rule out this interpretation but rather suggest that cebranopadol reduces nonspecific arousal linked to drug seeking that is often observed in yohimbine stress as well as footshock stress.

Cebranopadol is a pan-opioid agonist that binds with high affinity at MOP and NOP receptors. At higher concentrations it also activates KOP and DOP receptors (25). The pharmacological profile of cebranopadol partially overlaps with that of buprenorphine, a well characterized compound that acts as a high affinity partial MOP agonist and low affinity NOP agonist, but is an antagonist at KOP and DOP receptors (44). A wealth of evidence and several metanalyses support significant efficacy of buprenorphine in OUD. For instance, earlier studies showed that, compared to placebo, buprenorphine is more efficacious in maintaining patients free from illicit opioid use (45, 46). These data were confirmed by a robust meta-analysis of 21 double blind randomized clinical trials, in which buprenorphine was given for at least 3 weeks. Outcome measures were retention in buprenorphine treatment and illicit opioid use. The study found that buprenorphine is highly efficacious in maintaining patients free from illicit opioids use with better retention in treatment at higher buprenorphine doses (16-32 mg per day) compared with lower doses (47). On the other hand, studies in which long term retention in treatment was analyzed reported that after 6-months, the dropout rate was about 50%, indicating moderate efficacy for long-term relapse prevention (48). Another limitation of buprenorphine is its abuse potential and difficulty in discontinuing the treatment in patients that want to transition into a medication free state. For instance, a recent study in prescription opioid abusers showed that, compared to a standard maintenance therapy with a fixed buprenorphine dose, drug tapering led to a poorer clinical outcome with patients showing more days per week of illicit opioid use, fewer maximum consecutive weeks of opioid abstinence, and less retention in treatment. Ultimately, due to relapse, a large percentage of patient in the taper group had to be reintroduced to buprenorphine after weaning off the medication (49). These findings confirm that in OUD maintenance treatment, tapering remains a major challenge (50). In a recent rat place conditioning experiment, cebranopadol produced significant expression of preference for a cebranopalol-paired compartment, which is reflective of the drug’s ability to stimulate reward processes (29). On the other hand, we have demonstrated substantially lower expression of place preference in animals treated with cebranopadol, compared to those receiving morphine (30). Moreover, Tzschentke and colleagues (51) treated rats and mice with chronic cebranopadol to measure the development of physical dependence and expression of opioid-like withdrawal upon drug discontinuation. In that study, animals receiving cebranopadol demonstrated fewer signs of withdrawal compared to morphine. Most importantly, in our present evaluation of cebranopadol’s abuse potential, we found that whereas cebranopadol maintains operant responding under a FR1 contingency, reflecting positive motivational effects, it does not support operant self-administration under a PR contingency, which indicates that the motivation for the drug is low. This finding is indirectly supported by data from a phase II clinical trial in which lower back pain patients treated chronically with cebranopadol showed a good tolerability profile and no major signs of dependence (52).

Reinstatement experiments in rodents demonstrated that buprenorphine is not efficacious in attenuating heroin and cocaine seeking elicited by stress (6). Similar findings were reported with methadone (53). Notably, in buprenorphine-maintained heroin dependent patients, yohimbine increased opioid seeking (40). This indicates that medications currently approved for opioid maintenance therapies may have limited efficacy for stress-induced relapse prevention. In this respect cebranopadol can be expected to be superior since it reduced yohimbine stress-induced reinstatement of heroin seeking.

We hypothesize that the broader efficacy of cebranopadol depends upon its ability to activate the N/OFQ/NOP system. Activation of NOP prevents several actions of classical opioid agonists including tolerance to opioid-induced supraspinal analgesia (54), morphine-induced conditioned place preference (21, 55) and morphine induced increases in extracellular dopamine levels in the nucleus accumbens (56). Moreover, activation of NOP by N/OFQ or by synthetic agonists produces anxiolytic-like effects (57, 58) that appear to be particularly robust under stressful conditions, such as during drug withdrawal (59). This effect possibly depends on the ability of NOP agonists to act as functional antagonists for extra-hypothalamic actions of the corticotrophin releasing factor (CRF) and CRF1 receptor (CRF1R) (60–62). Antagonism or genetic deletion of CRF1R attenuates stress-induced reinstatement of morphine (63) and heroin seeking (64) and reduces the expression of somatic and affective opioid withdrawal signs (65–68). Moreover, results from a recent clinical study showed that pexacerfont, a selective CRF1R antagonist attenuates withdrawal symptoms in subjects with heroin/methamphetamine dependence (69). Cebranopadol, by activating NOP receptors, may indirectly antagonize CRF1R mediated actions during drug abstinence which may explain its ability to attenuate stress-induced relapse.

The role of NOP mechanisms in mediating cebranopadol effects was demonstrated by an earlier study showing that the compound attenuates cocaine self-administration through co-activation of NOP and MOP receptors (30).

Based on the present findings one may speculate that, compared to already approved medications for maintenance therapy in OUD (i.e., buprenorphine, methadone etc), cebranopadol may offer advantages such as lower abuse liability, broader efficacy for relapse prevention and possibly faster tapering in patients that are motivated to transition into a medication free state. Despite significant efforts over recent years to develop new medications for the treatment of OUD, very little innovation has been achieved. The rather advanced clinical development of cebranopadol makes this compound an ideal candidate for immediate clinical investigation in heroin-dependent patients.

## Aknowledgements

Authors wish to thank Agostino Marchi, Rina Righi and Alfredo Fiorelli for animal care and technical support.

## Authors and Contributions

VL, HL, QS, and FB run self-administration experiments. NC analysed data. FW, RN, LS, and NC contributed to design the experiments and provided inputs. NC and RC wrote the manuscript.

